# Association of age and colostrum discarding with exclusive breastfeeding in Ethiopia: systematic review and meta-analyses

**DOI:** 10.1101/405225

**Authors:** Sisay Mulugeta Alemu, Yihun Mulugeta Alemu, Tesfa Dejenie Habtewold

**Author notes:** Correspondence should be addressed to Sisay Mulugeta Alemu.

## Abstract

**Introduction:** Even though optimal breastfeeding is important, significantly low percentage of mothers’ initiate breastfeeding timely and maintain exclusive breastfeeding for 6 months. The aim of this meta-analyses and systematic review was to investigate whether maternal/caregivers’ age, infant age (0-6 months) and discarding colostrum affects timely initiation of breastfeeding (TIBF) and exclusive breastfeeding (EBF) in Ethiopia.

**Methods:** A systematic search of PubMed, SCOPUS, EMBASE, CINHAL, Web of Science and WHO Global Health Library electronic databases was done for all English published articles from 2000 to January 2018, supplemented by manual search of identified articles and grey literatures bibliographies. Two reviewers independently screened, extracted and graded the quality studies using Newcastle–Ottawa Scale (NOS). Heterogeneity was assessed using the I^2^ and Cochran Chi-square statistics. A weighted inverse variance random-effects model meta-analysis was done.

**Result:** A total of 37 articles (i.e., 14 studies on TIBF and 23 on EBF) were included. TIBF was associated with colostrum discarding (Odds ratio (OR) = 0.38, 95% CI = 0.21-0.68) but not with maternal/caregivers’ age (OR = 0.98, 95% CI = 0.83-1.15). In addition, colostrum discarding (OR = 0.56, 95% CI = 0.37-0.84) and infant age (OR = 1.86, 95% CI = 1.45-2.39) were significantly associated with EBF but not maternal/caregivers’ age (OR = 1.07, 95% CI = 0.81-1.40).

**Conclusion:** This meta-analyses indicated absence of association between maternal/caregivers’ age and breastfeeding practice. Colostrum discarding was associated with both EBF and TIBE. This evidence could be helpful to counsel all reproductive age mothers and who discard colostrum.

## Introduction

World Health Organization (WHO) and United Nation Children’s Fund (UNICEF) defines timely initiation of breastfeeding (TIBF) as putting a new born to breast within one hour of birth and exclusive breastfeeding (EBF) as feeding infants only human milk through breastfeeding or expressed breast milk and no other liquids or solids, except for drops or syrups with nutritional supplements or medicine.^(1)^ All infants should receive human within the first hour of birth, exclusively breastfed for the first six months and thereafter, nutritionally adequate and safe complementary foods to be introduced with continued breastfeeding for at least two years.^(2, 3)^ Breastfeeding is one of the smartest investment that prevents maternal and newborn morbidity and mortality.^(4-7)^ For example, TIBF and EBF prevents 22% and 60% of neonatal deaths respectively.^(2, 8, 9)^ Furthermore, exclusive breastfeeding for a longer duration benefits child neurodevelopment and increase IQ.^(10)^

Despite the aforementioned advantages, significantly low percentage of mothers initiate breastfeeding within the first hour of birth and maintain exclusive breastfeeding for 6 months. Globally, 44% and 40% of newborns breastfed with in the first hour and exclusively breastfeed for six months respectively.^(4, 11)^ In developing countries, the prevalence of TIBF ranges 22.4 to 52.8% ^(17-23)^ and EBF ranges 10 to 49.1%.^(11, 17, 18, 27, 28)^ In Ethiopia, based on our previous meta-analyses, (unpublished results) the national prevalence of TIBF and EBF is 67.5% and 60.5% respectively.

Previous studies have identified several associated factors, including maternal/caregiver’s age, newborn age and colostrum discarding, of timely TIBF and EBF.^(12-22)^ Previous studies shows that infant age and colostrum discarding have been associated with late initiation of breastfeeding and nonexclusive breastfeeding.^(18, 20, 23-25)^ Regarding maternal/caregiver’s age, most of the reviewed literatures reveals that older mothers practice TIBF^(14, 19, 20, 26)^ and EBF^(12, 17, 22, 27, 28)^ higher than young mothers although the age cut-off value varies between studies.

Another study,^(13)^ which measured age as a continuous variable, also concludes that increased maternal age positively associated TIBF and EBF. On the contrary, some studies showed that increased maternal age was associated with delayed initiation of breastfeeding and nonexclusive breastfeeding.^(15, 16)^ Furthermore, other studies showed absence of association.^(21, 29)^ Taken together, inconsistencies persisted and the association is inconclusive.

Hence, there is an urgent need to synthesize individual studies data to make a better conclusion on the association between maternal age, infant age and colostrum discarding and breastfeeding practice (i.e., TIBF and EBF). So far, several systematic reviews and meta-analyses have been conducted on TIBF and EBF.^(14, 18, 20, 30-32)^ In Ethiopia, only one meta-analysis investigated the association of place of residence and delivery with TIBF.^(32)^ In our previous meta-analyses, (unpublished results) we studied the association between maternal employment, breastfeeding counseling, model of delivery, place of delivery, sex of newborn, antenatal care and postnatal care and breastfeeding practice. We also investigated the association between TIBF and EBF. The present meta-analyses and systematic review aimed to determine whether maternal/caregivers age, infant age and colostrum discharging affects TIBF and EBF in Ethiopia. We hypothesized (i) increased maternal age positively associated with breastfeeding practice due to accumulated experience, (ii) increased infant age negatively associated with exclusive breastfeeding and (iii) colostrum discarding negatively associated with breastfeeding practice.

Following international recommendations, the Ethiopian government has taken steps to improving infant and young child feeding practices. Several national nutritional strategies,^(33)^ guidelines^(34)^ and nutrition programs^(35, 36)^ have been developed by Ministry of Health of Ethiopia since 2004. Likewise, the Health Sector Transformation Plan of Ethiopia^(37)^ has targeted to increase exclusive breast feeding to 72 % by 2020. Furthermore, Ethiopia has recently started celebrating world breastfeeding week every year.^(38)^ However, TIBF and EBF coverage is still below the WHO recommendation and attributed to several factors. This meta-analyses information could be valuable to provide updated evidence to develop national guidelines and strategies.

## Methods

### Protocol registration and publication

The protocol has been registered with the University of York Centre for Reviews and Dissemination International prospective register of systematic reviews (PROSPERO) (http://www.crd.york.ac.uk/PROSPERO/display_record.asp?ID=CRD42017056768) and published. ^(39)^

### Data source and search strategy

For all available publications, systematic search of PubMed, SCOPUS, EMBASE, Cumulative Index to Nursing and Allied Health Literature (CINAHL), Web of Science and WHO Global Health Library electronic databases was done. In addition, bibliographies of identified articles and grey literatures were hand-searched. A comprehensive search strategy was developed for each database in consultation with a medical information specialist (Supplementary file 1).

### Eligibility criteria

All studies published in English from 2000 to January 2018 were included. In addition, observational studies (cross-sectional, case–control, cohort, survey and surveillance reports) conducted in Ethiopia were included. However, studies on preterm newborn infants, infants in neonatal intensive care unit or a special care baby unit, low birth weight and mothers or infants with medical problems were excluded. Further, commentaries, anonymous reports, letters, duplicate studies, editorials, qualitative studies and citations without full text were excluded.

### Study screening and selection

All studies obtained from databases and manual search were exported to EndNote citation manager. The title and abstract of all studies were screened by reviewers (SM & TD) independently. Agreement between the reviewers, as measured by Cohen’s Kappa, was 0.76. Any disagreement was resolved by discussion. When consensus could not be reached, a third reviewer approved the final list of retained studies. A full-text review was performed by two independent investigators (SM & TD).

### Quality assessment and data extraction

Newcastle-Ottawa Scale (NOS), which has good inter-rater reliability and validity, was used to assess the quality of studies and for potential publication bias.^(40, 41)^ In addition, to define outcome measurements, WHO infant and young child feeding practice guideline was strictly followed. Based on previous systematic review report,^(14, 42, 43)^ maternal/Caregiver’s age was dichotomized as ≥ 25 versus <25 years old whereas infant age was dichotomized as ≤ 3 versus 3 to 6 months age. Joanna Briggs Institute (JBI) tool was used to extract the following data: study area (region and place), method (design), population, number of mothers (calculated sample size and participated in actual study) and cross-tabulated data. Geographic regions were categorized based on the current Federal Democratic Republic of Ethiopia administrative structure (Supplementary file 2). Discrepancies were resolved by consensus and cross-checking with the full-text.

### Statistical analysis

A weighted inverse variance random-effects model meta-analyses was implemented. Publication bias was assessed by visual inspection of funnel plot and Egger’s regression test for funnel plot asymmetry using standard error as a predictor in mixed-effects meta-regression model at p-value threshold ≤ 0.01.^(44)^ Duval and Tweedie trim-and-fill method^(45)^ was used if we found asymmetric funnel which indicate publication bias. Heterogeneity was assessed by Cochran’s Q X^2^ test (p-value ≤ 0.05) and I^2^ statistics (reference value > 80%).^(39)^ The data was analyzed using “metaphor” packages in R software version 3.2.1 for Window.

### Data synthesis and reporting

We analyzed the data in two groups of outcome measurements: TIBF and EBF. Results for each variable were shown using forest plots. Preferred Reporting Items for Systematic Reviews and Meta-Analyses (PRISMA) guideline was strictly followed (Supplementary file 3).

### Minor post hoc protocol changes

Before analysis was done, we made the following changes to our methods from the published protocol. We added the Joanna Briggs Institute (JBI) tool to extract the data. In addition, we used Duval and Tweedie trim-and-fill method to manage publication bias.

## Result

### Search results

We obtained 169 articles from PubMed, 24 from EMBASE, 200 from Web of Science, 85 from SCOPUS and 5 from other (CINHAL and WHO Global Health Library) electronic database searching. Forty-eight additional articles were found through a manual search of reference lists of included articles. After removing duplicates and screening of titles and abstracts, full-text of 82 studies were reviewed to assess eligibility. Forty-five articles were excluded after a full-text review due to several reasons: 19 studies on complementary feeding, 3 on pre-lacteal feeding, 3 on malnutrition, 19 with different variables of interest and one project review report. As a result, 37 articles (i.e., 14 studies on timely initiation of breastfeeding and 23 on exclusive breastfeeding) fulfilled the inclusion criteria and were included in the meta-analyses. The PRISMA flow diagram of literature screening and selection process is shown in figure 1.

**Figure 1:**
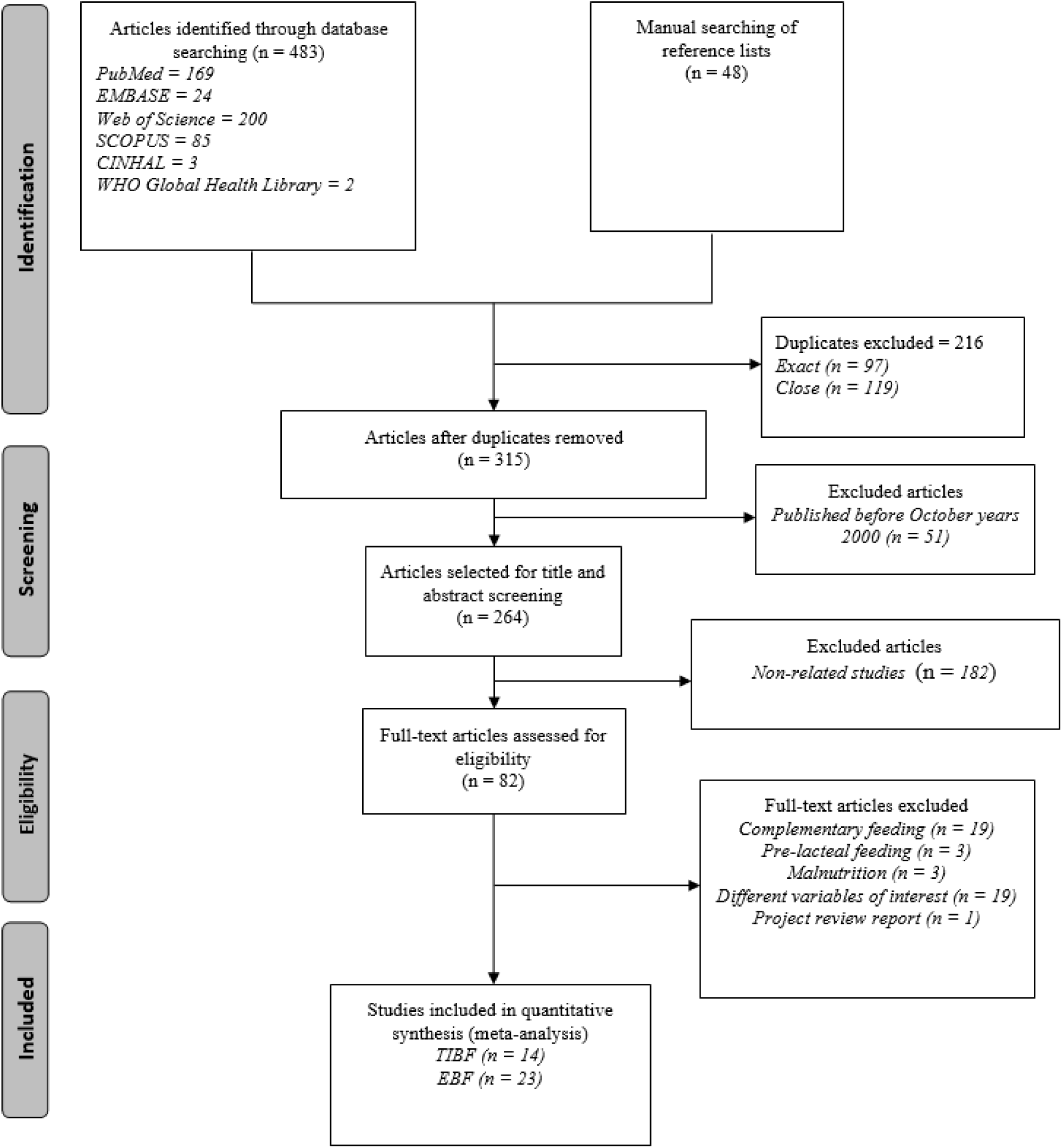
PRISMA flow diagram of literature screening and selection process; “n” in each stage represents the total number of studies that fulfilled a particular criterion.

### Study characteristics

Of these 14 studies on TIBF, most studies were conducted in Southern Nations Nationalities and People (SNNPR) region and Oromia region. Regarding maternal/caregiver’s residence, six and three studies conducted in urban and rural dwellers respectively (Table 1).

**Table 1:** Characteristics of studies included on TIBF

| Au-<br>thor/publicati<br>on year | Study area<br>(Region and<br>place) | Study meth-<br>od/desig<br>n | Study population | Calculated Sample<br>size/Participated | Factors | Breastfeeding initiation<br>(outcome) |  |  |
| --- | --- | --- | --- | --- | --- | --- | --- | --- |
|  |  |  |  |  |  | Within<br>1 hour | After 1<br>hour | Total |
| Maternal/Caregiver’s age versus timely initiation of breastfeeding |  |  |  |  |  |  |  |  |
| Wolde et.al.<br>2014 | Oromia,<br>Nekemte twon | Cross-sectional | Mothers who had<br>child less than 24<br>month | 182/174 | <25 years | 43 | 5 | 48 |
|  |  |  |  |  | ≥25 years | 111 | 15 | 126 |
|  |  |  |  |  | Total | 154 | 20 | 174 |
| Woldemichael<br>et.al. 2016 | Oromia, Tiyo<br>Woreda | Cross-sectional<br>study | mothers who have<br>children Less Than<br>One Year Age | 386/373 | <25 years | 83 | 39 | 122 |
|  |  |  |  |  | ≥25 years | 168 | 83 | 251 |
|  |  |  |  |  | Total | 251 | 122 | 373 |
| Adugna et.al<br>2014 | SNNPR, Arba<br>Minch Zuria | cross-sectional<br>study | Women who had<br>children under two<br>years | 384/383 | <25 years | 181 | 132 | 313 |
|  |  |  |  |  | ≥25 years | 38 | 32 | 70 |
|  |  |  |  |  | Total | 219 | 164 | 383 |
| Beyene et.al<br>2017 | SNNPR, Dale<br>Woreda | Cross-sectional<br>study | mothers of children<br>under 24 months | 634/ 634 | <25 years | 180 | 49 | 229 |
|  |  |  |  |  | ≥25 years | 337 | 52 | 389 |
|  |  |  |  |  | Total | 517 | 101 | 618 |
| Alemayehu<br>et.al 2014 | Tigray, Axum<br>town | cross<br>sectional<br>study | mothers who had<br>children aged 6-12<br>months | 418/418 | <25 years | 75 | 49 | 124 |
|  |  |  |  |  | ≥25 years | 169 | 125 | 294 |
|  |  |  |  |  | Total | 244 | 174 | 418 |
| Berhe et.al<br>2013 | Tigray,<br>Mekelle town | Cross-sectional<br>study | mothers of children<br>aged 0 to 24 months | 361/361 | <25 years | 120 | 27 | 147 |
|  |  |  |  |  | ≥25 years | 158 | 52 | 210 |
|  |  |  |  |  | Total | 278 | 79 | 357 |
| Setegn et.al<br>2011 | Oromia, Goba<br>district | cross<br>sectional<br>study | mothers with children<br>(< 12 months | 668/ 608 | <25 years | 107 | 108 | 215 |
|  |  |  |  |  | ≥25 years | 207 | 177 | 384 |
|  |  |  |  |  | Total | 314 | 285 | 599 |
| Tamiru et.al<br>2015 | SNNPR, Arba<br>Minch Zuria<br>Woreda | cross-sectional<br>study | mothers of infants<br>aged two years and<br>younger | 384/384 | <25 years | 150 | 109 | 259 |
|  |  |  |  |  | ≥25 years | 70 | 54 | 124 |
|  |  |  |  |  | Total | 220 | 163 | 383 |
| Regassa 2014 | SNNPR,<br>Sidama zone | Cross-sectional<br>study | with infants aged be-<br>tween 0 and 6<br>months old | 1100/ 1094 | <25 years | 354 | 77 | 431 |
|  |  |  |  |  | ≥25 years | 522 | 141 | 663 |
|  |  |  |  |  | Total | 876 | 218 | 1094 |
| Ekubay et al<br>2018 | Addis Ababa<br>Town | Cross-sectional<br>study | mothers with infants<br>younger than or equal<br>to six months of age | 597/583 | <25 years | 134 | 94 | 228 |
|  |  |  |  |  | ≥25 years | 195 | 141 | 336 |
|  |  |  |  |  | Total | 329 | 235 | 564 |
| Discarding colostrum versus timely initiation of breastfeeding |  |  |  |  |  |  |  |  |
| Wolde et.al.<br>2014 | Oromia,<br>Nekemte town | Cross-sectional<br>study | mothers who had<br>child less than 24<br>month | 182/174 | Discarding | 10 | 3 | 13 |
|  |  |  |  |  | No | 144 | 17 | 161 |
|  |  |  |  |  | Total | 154 | 20 | 174 |
| Adugna et.al.<br>2014 | SNNPR,<br>Hawassa city | Cross-sectional | Mothers with infants<br>aged 0–6 months | 541/529 | Discarding | 21 | 21 | 42 |
|  |  |  |  |  | No | 198 | 143 | 341 |
|  |  | study |  |  | Total | 219 | 164 | 383 |
| Hailemariam et.al. 2015 | Oromia, East Wollega zone | cross-sectional study | Mothers who had children less than 24 months | 594/593 | Discarding | 30 | 15 | 45 |
|  |  |  |  |  | No | 443 | 81 | 524 |
|  |  |  |  |  | Total | 473 | 96 | 569 |
| Tewabe 2016 | Amhara, Mot-ta town | cross-sectional study | mothers with infant less than six month old | 423/405 | Discarding | 49 | 33 | 82 |
|  |  |  |  |  | No | 270 | 53 | 323 |
|  |  |  |  |  | Total | 319 | 86 | 405 |
| Tilahun et.al. 2016 | Amhara, Debre Berhan town | cross sectional study | mothers who had children less than six months of age | 416/ 409 | Discarding | 15 | 46 | 61 |
|  |  |  |  |  | No | 241 | 91 | 332 |
|  |  |  |  |  | Total | 256 | 137 | 393 |
| Liben and Yesuf 2016 | Afar, Amibara district | Cross-sectional study | mothers of children aged less than 24 months | 407/ 403 | Discarding | 83 | 142 | 225 |
|  |  |  |  |  | No | 68 | 88 | 156 |
|  |  |  |  |  | Total | 151 | 230 | 381 |

Majority of studies on EBF were done in Amhara, SNNP and Oromia regions with 7, 6 and 3 studies respectively. Likewise, nine and seven studies conducted in urban and rural dwellers respectively. Furthermore, two studies used a nationally representative data of Ethiopian Demographic and Health Survey (EDHS) (Table 2).

**Table 2:**
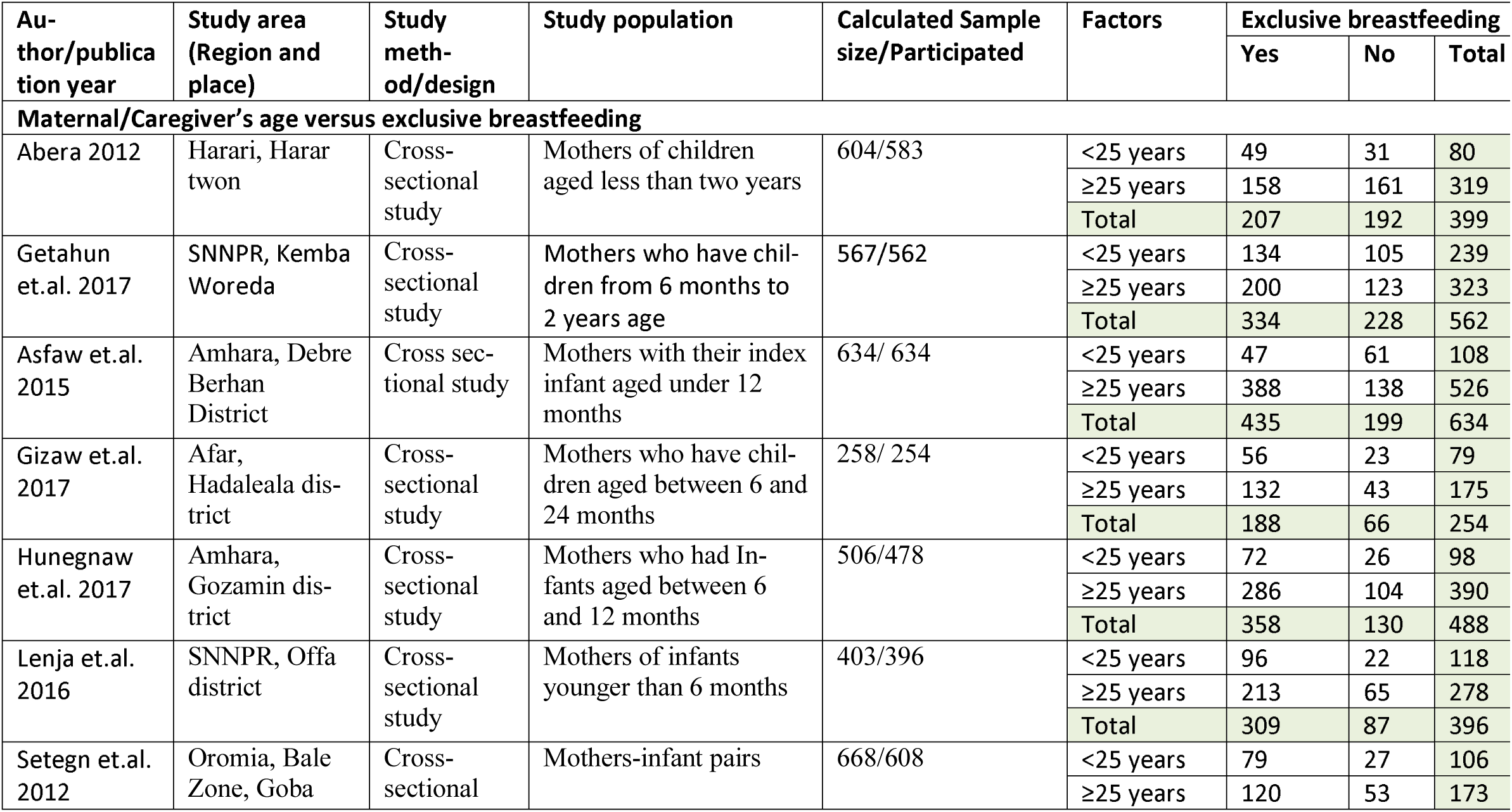
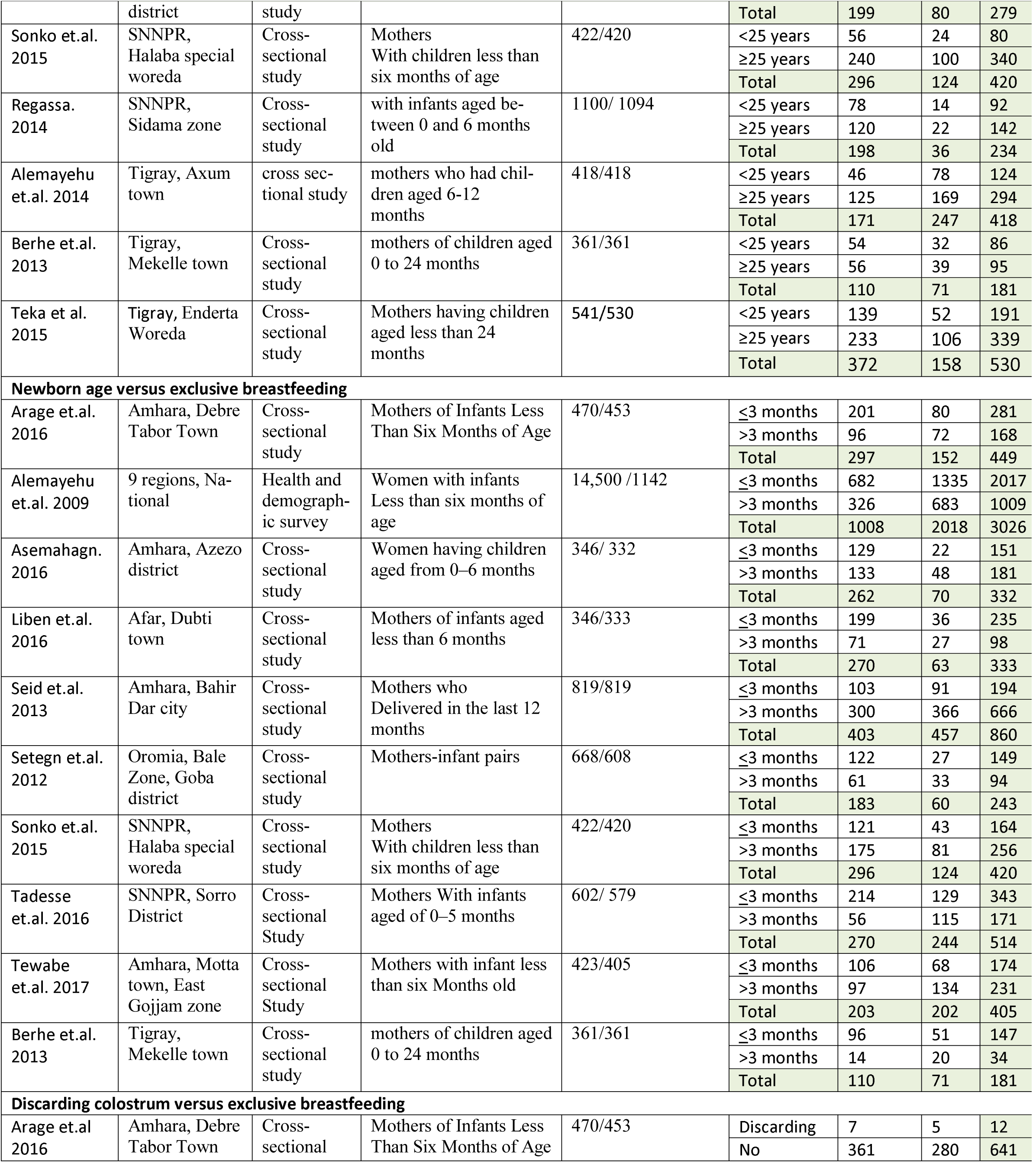
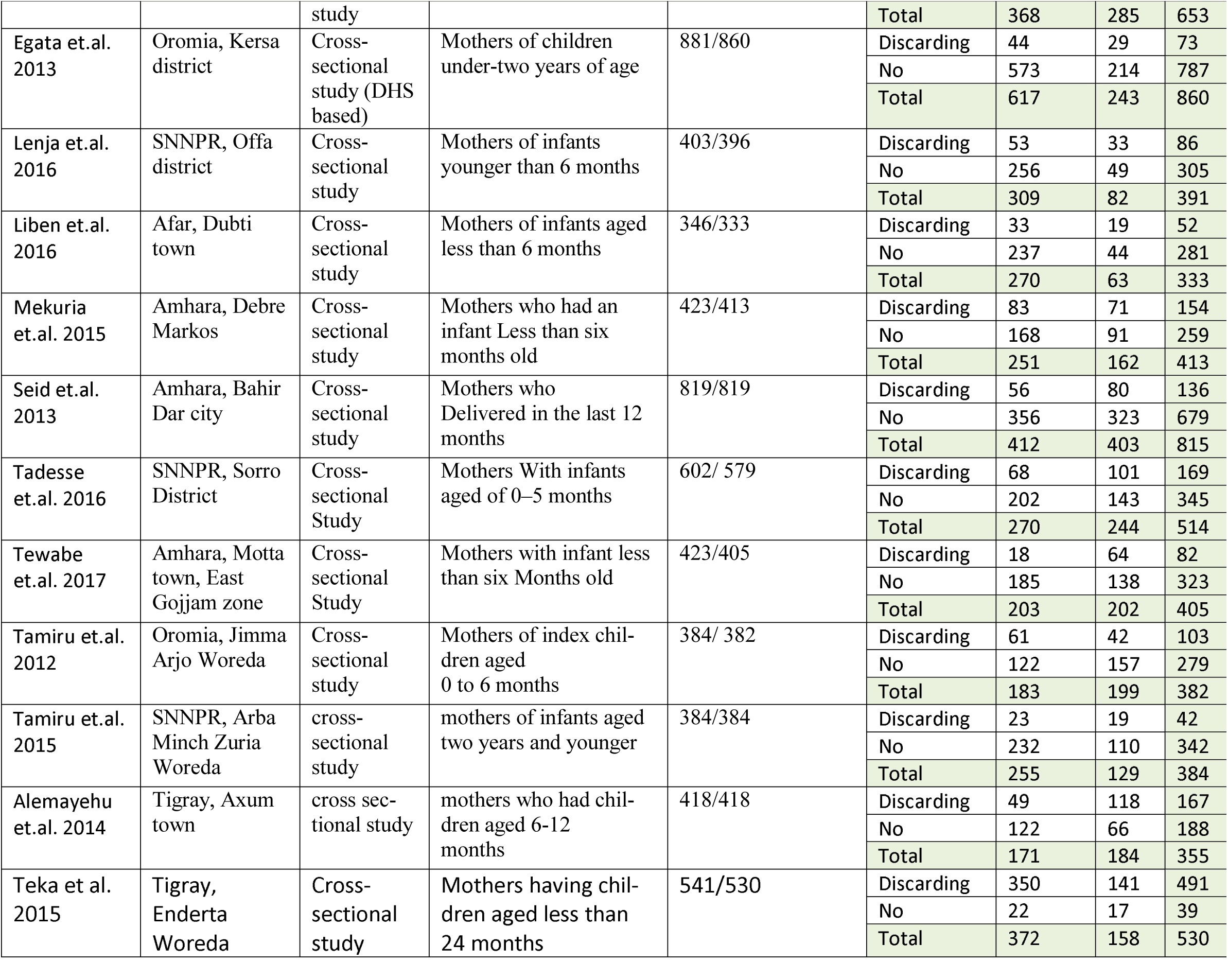
Characteristics of studies included on EBF

| Au-<br>thor/publica<br>tion year | Study area<br>(Region and<br>place) | Study meth-<br>od/design | Study population | Calculated Sample<br>size/Participated | Factors | Exclusive breastfeeding |  |  |
| --- | --- | --- | --- | --- | --- | --- | --- | --- |
|  |  |  |  |  |  | Yes | No | Total |
| Maternal/Caregiver's age versus exclusive breastfeeding |  |  |  |  |  |  |  |  |
| Abera 2012 | Harari, Harar<br>twon | Cross-sectional<br>study | Mothers of children<br>aged less than two years | 604/583 | <25 years | 49 | 31 | 80 |
|  |  |  |  |  | ≥25 years | 158 | 161 | 319 |
|  |  |  |  |  | Total | 207 | 192 | 399 |
| Getahun<br>et.al. 2017 | SNNPR, Kemba<br>Woreda | Cross-sectional<br>study | Mothers who have chil-<br>dren from 6 months to<br>2 years age | 567/562 | <25 years | 134 | 105 | 239 |
|  |  |  |  |  | ≥25 years | 200 | 123 | 323 |
|  |  |  |  |  | Total | 334 | 228 | 562 |
| Asfaw et.al.<br>2015 | Amhara, Debre<br>Berhan<br>District | Cross sec-<br>tional study | Mothers with their index<br>infant aged under 12<br>months | 634/ 634 | <25 years | 47 | 61 | 108 |
|  |  |  |  |  | ≥25 years | 388 | 138 | 526 |
|  |  |  |  |  | Total | 435 | 199 | 634 |
| Gizaw et.al.<br>2017 | Afar,<br>Hadaleala dis-<br>trict | Cross-sectional<br>study | Mothers who have chil-<br>dren aged between 6 and<br>24 months | 258/ 254 | <25 years | 56 | 23 | 79 |
|  |  |  |  |  | ≥25 years | 132 | 43 | 175 |
|  |  |  |  |  | Total | 188 | 66 | 254 |
| Hunegnaw<br>et.al. 2017 | Amhara,<br>Gozamin dis-<br>trict | Cross-sectional<br>study | Mothers who had In-<br>fants aged between 6<br>and 12 months | 506/478 | <25 years | 72 | 26 | 98 |
|  |  |  |  |  | ≥25 years | 286 | 104 | 390 |
|  |  |  |  |  | Total | 358 | 130 | 488 |
| Lenja et.al.<br>2016 | SNNPR, Offa<br>district | Cross-sectional<br>study | Mothers of infants<br>younger than 6 months | 403/396 | <25 years | 96 | 22 | 118 |
|  |  |  |  |  | ≥25 years | 213 | 65 | 278 |
|  |  |  |  |  | Total | 309 | 87 | 396 |
| Setegn et.al.<br>2012 | Oromia, Bale<br>Zone, Goba | Cross-sectional | Mothers-infant pairs | 668/608 | <25 years | 79 | 27 | 106 |
|  |  |  |  |  | ≥25 years | 120 | 53 | 173 |
|  | district | study |  |  | Total | 199 | 80 | 279 |
| Sonko et.al. 2015 | SNNPR, Halaba special woreda | Cross-sectional study | Mothers With children less than six months of age | 422/420 | <25 years | 56 | 24 | 80 |
|  |  |  |  |  | ≥25 years | 240 | 100 | 340 |
|  |  |  |  |  | Total | 296 | 124 | 420 |
| Regassa. 2014 | SNNPR, Sidama zone | Cross-sectional study | with infants aged between 0 and 6 months old | 1100/ 1094 | <25 years | 78 | 14 | 92 |
|  |  |  |  |  | ≥25 years | 120 | 22 | 142 |
|  |  |  |  |  | Total | 198 | 36 | 234 |
| Alemayehu et.al. 2014 | Tigray, Axum town | cross sectional study | mothers who had children aged 6-12 months | 418/418 | <25 years | 46 | 78 | 124 |
|  |  |  |  |  | ≥25 years | 125 | 169 | 294 |
|  |  |  |  |  | Total | 171 | 247 | 418 |
| Berhe et.al. 2013 | Tigray, Mekelle town | Cross-sectional study | mothers of children aged 0 to 24 months | 361/361 | <25 years | 54 | 32 | 86 |
|  |  |  |  |  | ≥25 years | 56 | 39 | 95 |
|  |  |  |  |  | Total | 110 | 71 | 181 |
| Teka et al. 2015 | Tigray, Enderta Woreda | Cross-sectional study | Mothers having children aged less than 24 months | 541/530 | <25 years | 139 | 52 | 191 |
|  |  |  |  |  | ≥25 years | 233 | 106 | 339 |
|  |  |  |  |  | Total | 372 | 158 | 530 |
| Arage et.al. 2016 | Amhara, Debre Tabor Town | Cross-sectional study | Mothers of Infants Less Than Six Months of Age | 470/453 | ≤3 months | 201 | 80 | 281 |
|  |  |  |  |  | >3 months | 96 | 72 | 168 |
|  |  |  |  |  | Total | 297 | 152 | 449 |
| Alemayehu et.al. 2009 | 9 regions, National | Health and demographic survey | Women with infants Less than six months of age | 14,500 /1142 | ≤3 months | 682 | 1335 | 2017 |
|  |  |  |  |  | >3 months | 326 | 683 | 1009 |
|  |  |  |  |  | Total | 1008 | 2018 | 3026 |
| Asemahagn. 2016 | Amhara, Azezo district | Cross-sectional study | Women having children aged from 0–6 months | 346/ 332 | ≤3 months | 129 | 22 | 151 |
|  |  |  |  |  | >3 months | 133 | 48 | 181 |
|  |  |  |  |  | Total | 262 | 70 | 332 |
| Liben et.al. 2016 | Afar, Dubti town | Cross-sectional study | Mothers of infants aged less than 6 months | 346/333 | ≤3 months | 199 | 36 | 235 |
|  |  |  |  |  | >3 months | 71 | 27 | 98 |
|  |  |  |  |  | Total | 270 | 63 | 333 |
| Seid et.al. 2013 | Amhara, Bahir Dar city | Cross-sectional study | Mothers who Delivered in the last 12 months | 819/819 | ≤3 months | 103 | 91 | 194 |
|  |  |  |  |  | >3 months | 300 | 366 | 666 |
|  |  |  |  |  | Total | 403 | 457 | 860 |
| Setegn et.al. 2012 | Oromia, Bale Zone, Goba district | Cross-sectional study | Mothers-infant pairs | 668/608 | ≤3 months | 122 | 27 | 149 |
|  |  |  |  |  | >3 months | 61 | 33 | 94 |
|  |  |  |  |  | Total | 183 | 60 | 243 |
| Sonko et.al. 2015 | SNNPR, Halaba special woreda | Cross-sectional study | Mothers With children less than six months of age | 422/420 | ≤3 months | 121 | 43 | 164 |
|  |  |  |  |  | >3 months | 175 | 81 | 256 |
|  |  |  |  |  | Total | 296 | 124 | 420 |
| Tadesse et.al. 2016 | SNNPR, Sorro District | Cross-sectional Study | Mothers With infants aged of 0–5 months | 602/ 579 | ≤3 months | 214 | 129 | 343 |
|  |  |  |  |  | >3 months | 56 | 115 | 171 |
|  |  |  |  |  | Total | 270 | 244 | 514 |
| Tewabe et.al. 2017 | Amhara, Motta town, East Gojjam zone | Cross-sectional Study | Mothers with infant less than six Months old | 423/405 | ≤3 months | 106 | 68 | 174 |
|  |  |  |  |  | >3 months | 97 | 134 | 231 |
|  |  |  |  |  | Total | 203 | 202 | 405 |
| Berhe et.al. 2013 | Tigray, Mekelle town | Cross-sectional study | mothers of children aged 0 to 24 months | 361/361 | ≤3 months | 96 | 51 | 147 |
|  |  |  |  |  | >3 months | 14 | 20 | 34 |
|  |  |  |  |  | Total | 110 | 71 | 181 |
| Arage et.al 2016 | Amhara, Debre Tabor Town | Cross-sectional | Mothers of Infants Less Than Six Months of Age | 470/453 | Discarding | 7 | 5 | 12 |
|  |  |  |  |  | No | 361 | 280 | 641 |
|  |  | study |  |  | Total | 368 | 285 | 653 |
| Egata et.al. 2013 | Oromia, Kersa district | Cross-sectional study (DHS based) | Mothers of children under-two years of age | 881/860 | Discarding | 44 | 29 | 73 |
|  |  |  |  |  | No | 573 | 214 | 787 |
|  |  |  |  |  | Total | 617 | 243 | 860 |
| Lenja et.al. 2016 | SNNPR, Offa district | Cross-sectional study | Mothers of infants younger than 6 months | 403/396 | Discarding | 53 | 33 | 86 |
|  |  |  |  |  | No | 256 | 49 | 305 |
|  |  |  |  |  | Total | 309 | 82 | 391 |
| Liben et.al. 2016 | Afar, Dubti town | Cross-sectional study | Mothers of infants aged less than 6 months | 346/333 | Discarding | 33 | 19 | 52 |
|  |  |  |  |  | No | 237 | 44 | 281 |
|  |  |  |  |  | Total | 270 | 63 | 333 |
| Mekuria et.al. 2015 | Amhara, Debre Markos | Cross-sectional study | Mothers who had an infant Less than six months old | 423/413 | Discarding | 83 | 71 | 154 |
|  |  |  |  |  | No | 168 | 91 | 259 |
|  |  |  |  |  | Total | 251 | 162 | 413 |
| Seid et.al. 2013 | Amhara, Bahir Dar city | Cross-sectional study | Mothers who Delivered in the last 12 months | 819/819 | Discarding | 56 | 80 | 136 |
|  |  |  |  |  | No | 356 | 323 | 679 |
|  |  |  |  |  | Total | 412 | 403 | 815 |
| Tadesse et.al. 2016 | SNNPR, Sorro District | Cross-sectional Study | Mothers With infants aged of 0–5 months | 602/ 579 | Discarding | 68 | 101 | 169 |
|  |  |  |  |  | No | 202 | 143 | 345 |
|  |  |  |  |  | Total | 270 | 244 | 514 |
| Tewabe et.al. 2017 | Amhara, Motta town, East Gojjam zone | Cross-sectional Study | Mothers with infant less than six Months old | 423/405 | Discarding | 18 | 64 | 82 |
|  |  |  |  |  | No | 185 | 138 | 323 |
|  |  |  |  |  | Total | 203 | 202 | 405 |
| Tamiru et.al. 2012 | Oromia, Jimma Arjo Woreda | Cross-sectional study | Mothers of index children aged 0 to 6 months | 384/ 382 | Discarding | 61 | 42 | 103 |
|  |  |  |  |  | No | 122 | 157 | 279 |
|  |  |  |  |  | Total | 183 | 199 | 382 |
| Tamiru et.al. 2015 | SNNPR, Arba Minch Zuria Woreda | cross-sectional study | mothers of infants aged two years and younger | 384/384 | Discarding | 23 | 19 | 42 |
|  |  |  |  |  | No | 232 | 110 | 342 |
|  |  |  |  |  | Total | 255 | 129 | 384 |
| Alemayehu et.al. 2014 | Tigray, Axum town | cross sectional study | mothers who had children aged 6-12 months | 418/418 | Discarding | 49 | 118 | 167 |
|  |  |  |  |  | No | 122 | 66 | 188 |
|  |  |  |  |  | Total | 171 | 184 | 355 |
| Teka et al. 2015 | Tigray, Enderta Woreda | Cross-sectional study | Mothers having children aged less than 24 months | 541/530 | Discarding | 350 | 141 | 491 |
|  |  |  |  |  | No | 22 | 17 | 39 |
|  |  |  |  |  | Total | 372 | 158 | 530 |

### Timely initiation of breastfeeding (TIBF)

Among 14 studies, 10 studies^(46-55)^ reported the association between TIBF and maternal/caregiver’s age in 4,963 mothers. The pooled odds ratio (OR) of maternal/caregiver’s age was 0.98 (95% CI 0.83 - 1.15, p = 0.78) (figure 2). Although not statistically significant, mothers ≥ 25 years old age had 2% lower chance of initiating breastfeeding within one hour compared to their younger counterparts. Egger’s regression test for funnel plot asymmetry was not significant (z = −0.40, p = 0.69).

**Figure 2:**
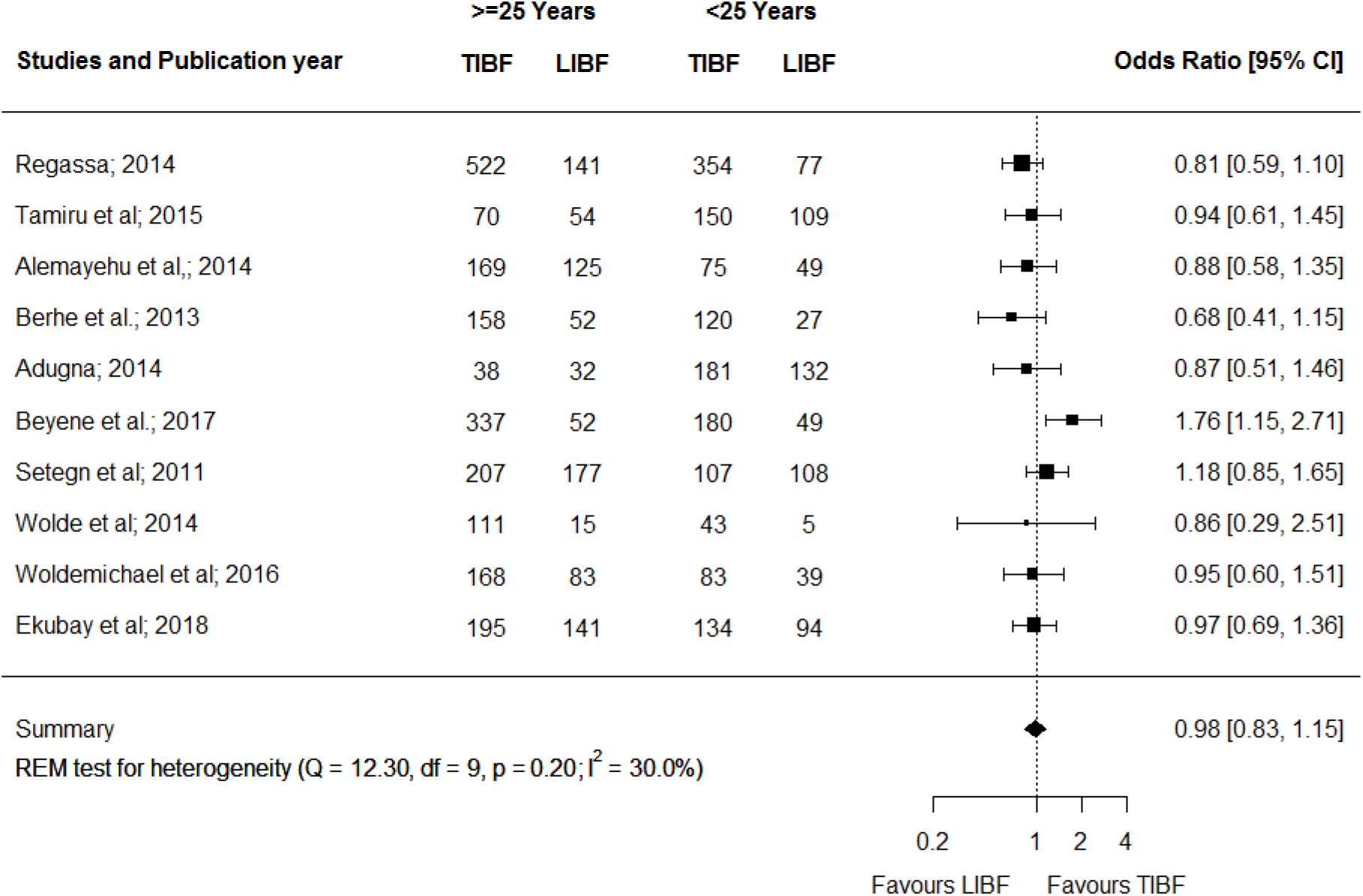
Forest plot of the unadjusted odds ratios with corresponding 95% CIs of studies on the association of maternal/caregiver’s age and TIBF. The horizontal line represents the confidence interval, the box and its size in the middle of the horizontal line represent the weight of sample size. The polygon represents the pooled odds ratio. TIBF = timely initiation of breastfeeding; LIBF = late initiation of breastfeeding; REM = random-effects model.

Likewise, 6 out of 14 studies reported the association between TIBF and colostrum discarding in 2,305 mothers ^(46, 48, 56-59)^. The pooled OR of colostrum discarding was found to be 0.38 (95% CI 0.21 – 0.68, p = 0.001) (figure 3). Compared to mothers who feed colostrum, mothers who discard colostrum had 62% significantly lower chance of initiating breastfeeding within one hour. Egger’s regression test for funnel plot asymmetry was not significant (z = −0.24, p = 0.81).

**Figure 3:**
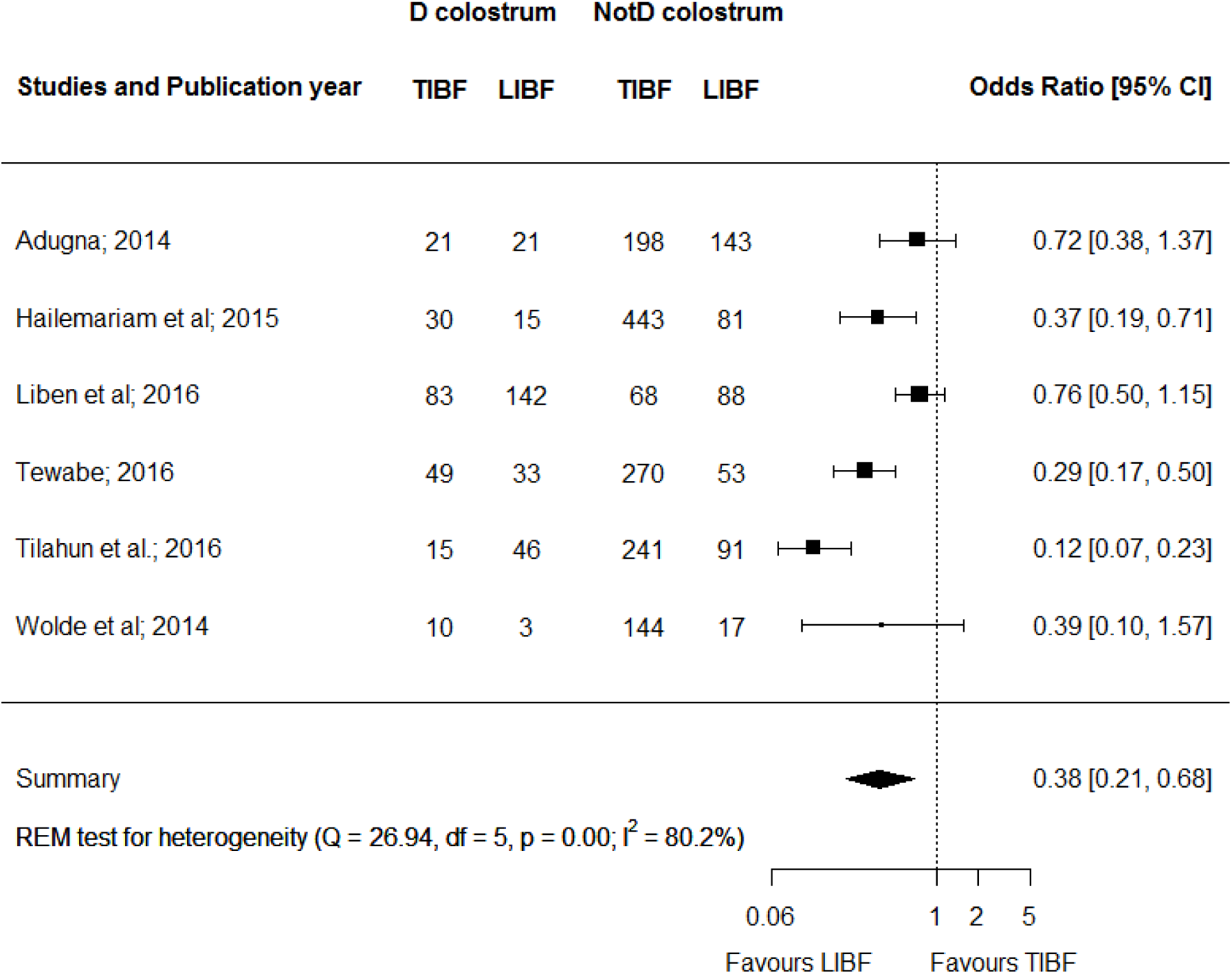
Forest plot of the unadjusted odds ratios with corresponding 95% CIs of studies on the association of colostrum discarding and TIBF. The horizontal line represents the confidence interval, the box and its size in the middle of the horizontal line represent the weight of sample size. The polygon represents the pooled odds ratio. TIBF = timely initiation of breastfeeding; LIBF = late initiation of breastfeeding; REM = random-effects model; D=Discarding; NotD = Not discarding.

### Exclusive breastfeeding

Twelve studies^(50, 51, 54, 60-68)^ involving 4,929 individuals reported the association between EBF and maternal/caregiver’s age. As showed in figure 4, the pooled OR of maternal/caregiver’s age was 1.07 (95% CI 0.81 - 1.40, p = 0.63). Mothers ≥ 25 years old age had 7% higher chance of exclusively breastfeeding during the first six months compared to mothers <25 years old; however, it was not statistically significant. Egger’s regression test for funnel plot asymmetry was not significant (z = −0.99, p = 0.32).

**Figure 4:**
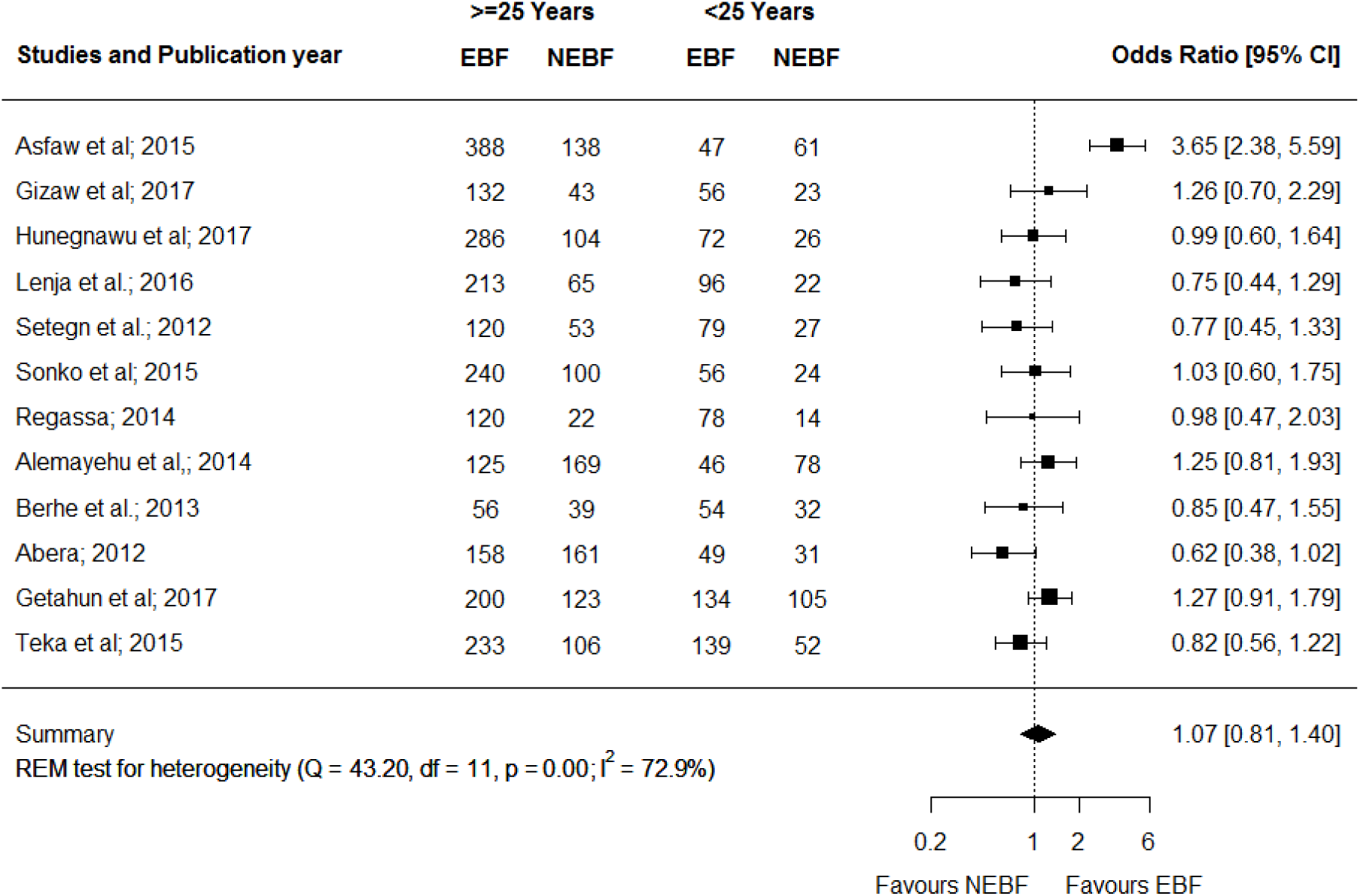
Forest plot of the unadjusted odds ratios with corresponding 95% CIs of studies on the association of maternal/caregiver’s age and EBF. The horizontal line represents the confidence interval, the box and its size in the middle of the horizontal line represent the weight of sample size. The polygon represents the pooled odds ratio. EBF = Exclusive breastfeeding; NEBF = Non-exclusive of breastfeeding; REM = random-effects model.

In addition, ten^(51, 65, 66, 69-75)^ out of 23 studies reported the association between EBF and infant age with a total sample of 6,763 mothers. The pooled OR of infant age was 1.86 (95% CI 1.45 - 2.39, p <0.001) (figure 5). Children ≤3 months old had 86% statistically significant higher chance of being exclusively breastfed compared to children older than 3 months. Egger’s regression test for funnel plot asymmetry was not significant (z = 2.31, p = 0.02).

**Figure 5:**
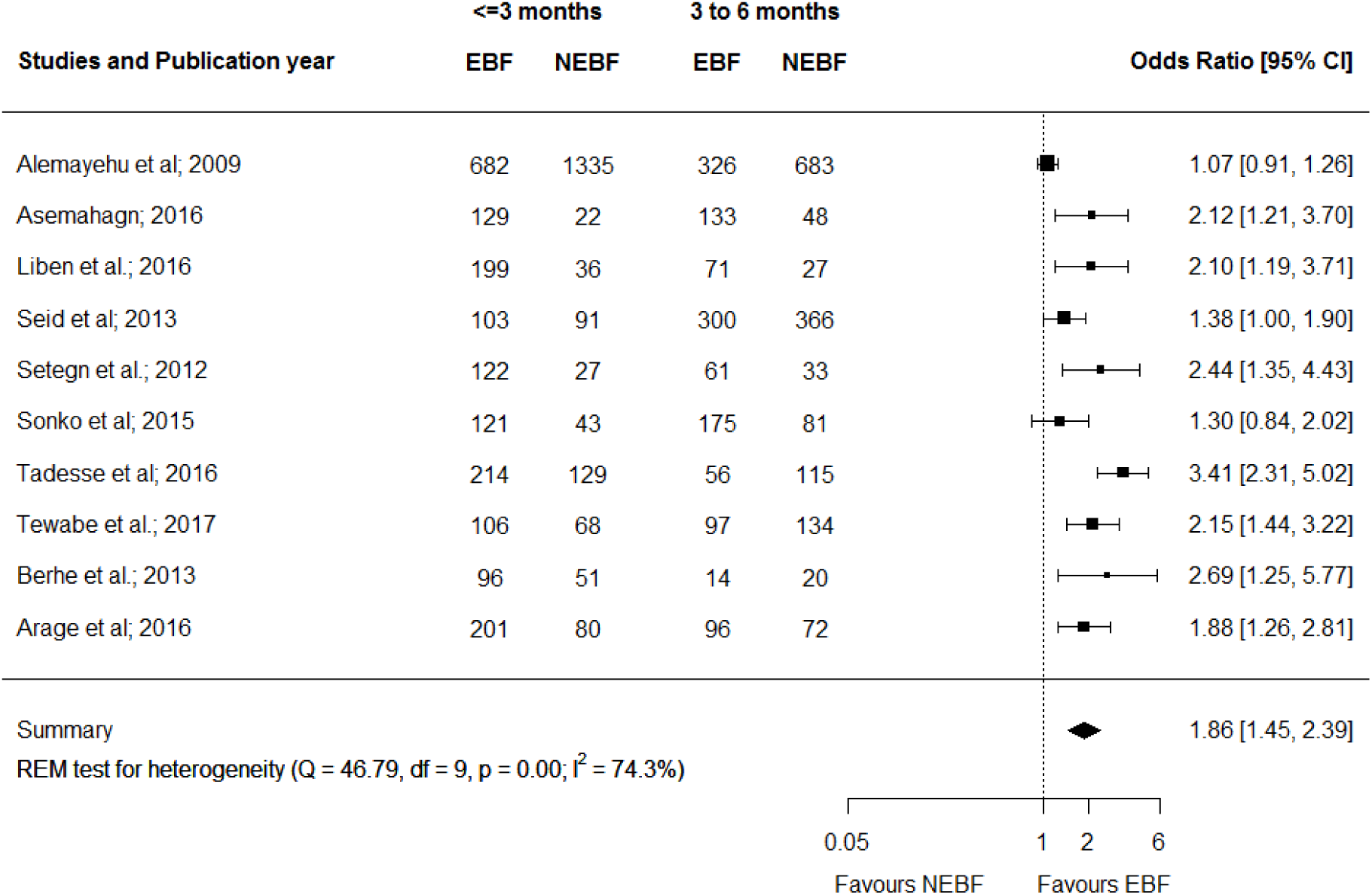
Forest plot of the unadjusted odds ratios with corresponding 95% CIs of studies on the association of infant age and EBF. The horizontal line represents the confidence interval, the box and its size in the middle of the horizontal line represent the weight of sample size. The polygon represents the pooled odds ratio. EBF=Exclusive breastfeeding; NEBF=Non-exclusive of breastfeeding; REM=random effects model.

Finally, 12 studies^(50, 53, 64, 68, 71-78)^ reported the association between EBF and colostrum discarding with a sample of 6,035 mothers. As indicated in figure 6, the pooled OR of colostrum discarding was 0.56 (95% CI 0.37 - 0.84, p = 0.005). Mothers who discard colostrum had 44% statistically significant lower chance of exclusively breastfeeding during the first 6 months compared to mothers who feed colostrum. Egger’s regression test for funnel plot asymmetry was not significant (z = 0.68, p = 0.49).

**Figure 6:**
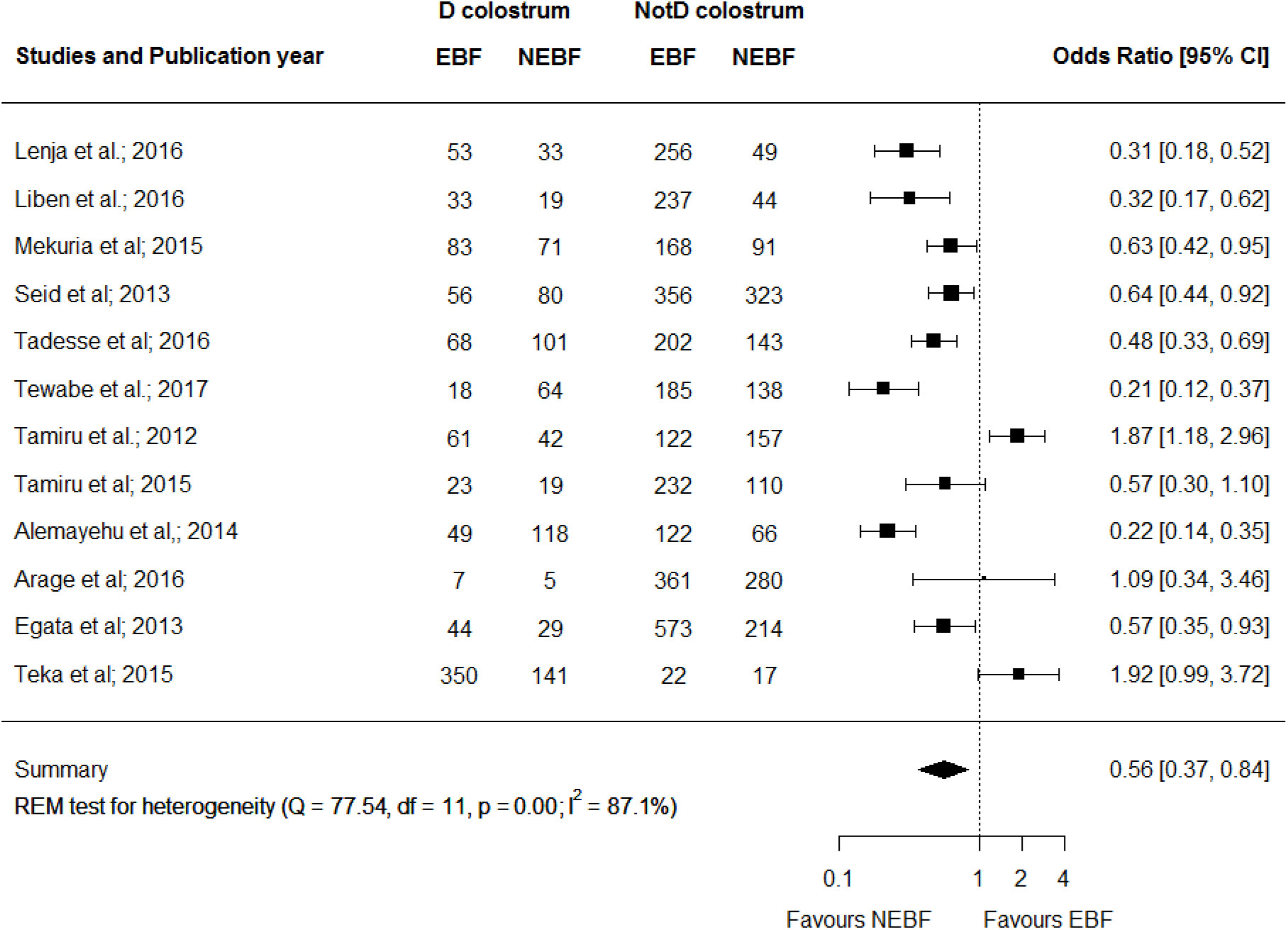
Forest plot of the unadjusted odds ratios with corresponding 95% CIs of studies on the association of discarding colostrum and EBF. The horizontal line represents the confidence interval, the box and its size in the middle of the horizontal line represent the weight of sample size. The polygon represents the pooled odds ratio. EBF = Exclusive breastfeeding; NEBF = Non-exclusive breastfeeding; REM = random-effects model; D = Discarding; NotD = Not discarding.

## Discussion

This study examined the associations of TIBF and EBF with colostrum discarding, maternal/caregiver’s age and infant age. To our knowledge, this is the first systematic review and meta-analyses in this topic to-date in Ethiopia. This meta-analysis uncovered colostrum discarding was significantly associated with TIBF but not maternal/caregiver’s age. On the other hand, colostrum discarding and infant age was found to be significantly associated with EBF but not maternal/caregiver’s age.

We found that colostrum discarding was significantly associated with TIBF. Mothers who discard colostrum had 62% significantly lower chance of initiating breastfeeding within one hour compared to mothers who feed colostrum to their child. This may be explained by the attempt of discard colostrum to get white milk may take time which therefore results in a delayed initiation of breastfeeding.

In the present meta-analysis, we found a statistically significant association between EBF and infant age. This finding confirmed our hypothesis and consistent with a large body of evidence showing that increased infant age is negatively associated with exclusive breastfeeding.^(18, 20, 23, 24, 79, 80)^ This may be due to the fact that giving traditional post-partum care and support is common in Ethiopia immediately after birth which may create opportunity for the mother to exclusively breastfeed the child. Since, this traditional post-partum care and support decreases as the age of the infant increases, it may lead the mother to work outside. This may therefore force the mother to stop EBF. Evidenced worldwide also agreed on the point that presence of social support is associated with better breastfeeding outcome.^(81-83)^ Another possible reason is the workload and short maternity leave in Ethiopia, which is only two months post-partum until recently, may influence the mother to withdraw EBF early. This hypothesis was supported by our previous meta-analyses whereby maternal employment significantly lower EBF and other studies.^(83, 84)^ Moreover, this could also be related to the short birth interval in Ethiopia.

We noted that colostrum discarding significantly associated with EBF. The finding was in line with studies conducted in Nepal ^(85)^ and Laos.^(86)^ This may be due to the fact that discarding colostrum leading to pre-lacteal feeding. In agreement with recent studies,^(87-92)^ maternal/caregiver’s age was not significantly associated with either EBF or TIBF. This is against our hypothesis and disproves the notion that older mothers have better breastfeeding experience than young mothers that helps them to practice optimal TIBF and EBF. However, there is robust evidence that supported all reproductive age group mothers can maintain optimal TIBF and EBF equally.^(24, 84, 93)^ Therefore, the discrepancy may be due to the following reasons: (1) most studies used maternal age rather than age at first birth; (2) different studies have used different age categories; and (3) breastfeeding is not age dependent or can be confounded by innate maternal behavior.

This meta-analyses study has several implications. It provided evidence on breastfeeding practice and its associated factors in an Ethiopian context, which can be useful for cross-country / cross-cultural comparison and for breastfeeding improvement initiative in Ethiopia. The present study provides an overview of up-to-date evidence for nutritionist and public health professionals. The findings also indicate emphasis should be given for all age group of mothers/caregivers during breastfeeding intervention. Furthermore, this study points out colostrum discarding and associated believes should be considered during designing breastfeeding interventions.

The association was estimated in large sample size and recent and nationally representative studies were included. In addition, this systematic review and meta-analysis was conducted based on a registered and published protocol, and guidelines for the Meta-analysis of Observational Studies in Epidemiology (MOOSE) was strictly followed. This study has also several limitations. First, some studies were excluded because of the difference in age category. Second, almost all included studies were observational which hinder causality inference. Third, even though we have used broad search strategies, the possibility of missing relevant studies cannot be fully exempted. Fourth, based on the conventional methods of statistical testing, a few analyses suffer from high levels of between-study heterogeneity. The course of heterogeneity was carefully explored and may be due to difference of study area; therefore, the result should be interpreted with caution.

In conclusion, colostrum discarding was a possible barrier for both TIBF and EBF. Additionally, increased infant age were found to be a risk factor for non EBF. However, maternal/caregiver’s age was not a determinant factor for both TIBF and EBF. Interventions targeted on increasing the rate of TIBF and EBF should give special focus on colostrum discarding and associated beliefs. In addition, future research should be required to identify other factors affecting duration of EBF in Ethiopia. Further investigation is also required to assess the effect of age at first birth.

## Ethics approval

Not applicable.

## Contributors

TD and SM conceived and designed the study. TD developed syntax for searching databases and analyzed the data. TD and SM wrote and revised the manuscript. All the authors read and have given the final approval.

## Patient Consent

Not required, because the review will not employ primary data collection.

